# TSS-Captur: A User-Friendly Characterization Pipeline for Transcribed but Unclassified RNA transcripts

**DOI:** 10.1101/2024.07.05.602221

**Authors:** Mathias Witte Paz, Thomas Vogel, Kay Nieselt

**Affiliations:** Institute for Bioinformatics and Medical Informatics, University of Tübingen, 72076, Tübingen, Germany

## Abstract

RNA-seq and its 5’-enrichment-based methods for prokaryotes have enabled the base-exact identification of transcription starting sites (TSSs) and have improved gene expression analysis. Computational methods analyze this experimental data to identify TSSs and classify them based on proximal annotated genes. While some TSSs cannot be classified at all (orphan TSSs), other TSSs are found on the reverse strand of known genes (antisense TSSs), but are not associated with the direct transcription of any known gene. Here, we introduce `TSS-Captur`, a novel pipeline, that uses computational approaches to characterize genomic regions starting from experimentally confirmed, but unclassified TSSs. By analyzing experimental TSS data, `TSS-Captur` characterizes unclassified signals, hence complementing prokaryotic genome annotation tools and enhancing the bacterial transcriptome understanding. `TSS-Captur` classifies extracted transcripts into coding or non-coding genes and predicts for each putative transcript its transcription termination site. For non-coding genes, the secondary structure is computed. Furthermore, putative promoter regions are analyzed to identify enriched motifs. An interactive report allows a seamless data exploration. We validated `TSS-Captur` with a *Campylobacter jejuni* dataset and characterized unlabeled non-coding RNAs in *Streptomyces coelicolor*. Besides its usage over the command-line, `TSS-Captur` is available as a web-application to enhance its user accessibility and explorative capabilities.

## 1 Introduction

In molecular biology and genetics, the identification of transcription start sites (TSSs) can provide valuable insights into the regulation of gene expression in prokaryotic organisms, aiding researchers in understanding the mechanisms controlling transcription initiation, such as the identification of nearby transcription factor binding sites [30]. The base-specific identification of TSSs enables researchers to better understand how genes are regulated and how they respond to different environmental conditions or stimuli. Differential RNA-seq (dRNA-seq) [29] and Cappable-seq [12] are enrichment methods for RNA-seq that allow the identification of TSSs in prokaryotes. These methods adapt the libraries to be sequenced for identifying the 5^*′*^-end of an expressed gene. Such experiments typically result in hundreds to thousands of TSSs that make a manual determination cumbersome. Computational methods have therefore been developed in the past years to facilitate the process of TSS identification.

`TSSpredator` [11] is one of these tools which determines the TSS signals from the experimental data. The TSSs are then associated with annotated genes depending on their proximity and strand. Many approaches for determining genomewide TSS maps in a large variety of bacteria have led to many insights. One study computed a wide transcriptome analysis for *Helicobacter pylori* and was able to provide an overview of regulatory mechanisms found in the 5^*′*^-UTR of annotated genes [3]. A more recent study reported a similar map for *Bacteroides thetaiotaomicron* providing more insight into its vast transcription regulation mechanisms and identifying over 269 potential non-coding RNA (ncRNA) candidates, such as the *GibS*, a trans-RNA gene regulating the metabolism of the bacteria [26].

In these and in many further studies, numerous TSS signals were not found close to any annotated genes. Hence, such TSSs are typically reported as orphans (oTSSs). Moreover, other TSSs are located on the antisense strand within the coordinates of an annotated gene, but with no annotated gene in the direct downstream region (thus typically called antisense TSSs, aTSSs). Genome annotation pipelines, such as `Bakta` [27], `Prokka` [28] or `PGAP` [32] focus on the identification of protein-coding RNA genes (mRNAs) and house-keeping RNA genes. Non-coding RNA genes are often overlooked, since their diversity of patterns make their computational prediction more difficult, especially in prokaryotes [31, 28]. Still, they play an important role in prokaryotes, such as *cis* or *trans* small RNAs [5]. We hypothesize that oTSS and aTSS signals are linked to such ncRNA genes, as well as other overlooked protein-coding genes. Hence, providing an explorative tool for the characterization of such sites would provide more insight in the prokaryotic transcriptome. For example, it could provide a general idea of their function (e.g. mRNA or ncRNA), an approximate length of their transcript as well as their transcription termination site (TTS), and analyzing other features, such as their putative secondary structure or enriched motifs found in the promoter region.

To enable the characterization of these classes of TSSs and their surrounding genomic regions in an explorative manner, we have developed `TSS-Captur`, a TSS-characterization pipeline for transcribed but unclassified (RNA)-transcripts. `TSS-Captur` combines the results of transcriptomic experiments with the underlying genomic sequence to characterize putative transcripts starting on orphan and antisense TSS signals predicted using `TSSpredator`. For the overall characterization, `TSS-Captur` integrates different tools. `TSS-Captur` combines two well-established prediction methods — one based on comparative genomics (`QRNA`) and one *ab initio* method (`CNIT`) — to determine whether a transcript beginning with an oTSS or aTSS originates from a protein-coding or a non-coding gene. This allows `TSS-Captur` to account for multiple aspects of the prediction and strengthen its findings. Furthermore, `TSS-Captur` combines TSS signals with a possible transcription termination site (TTS) that has been predicted based on the genomic sequence. Moreover, it enables the user to explore the thermodynamic characteristics (i.e., folding potential) of possible identified ncRNA genes by visualizing their computed secondary structure. Lastly, it facilitates the analysis of motifs within the regions upstream of each TSS signal for the identification of transcription factor binding sites or other regulatory elements. All these results are shown in an interactive report to provide both an overview of the predicted results and detailed information for each transcript. `TSS-Captur` is implemented as an interactive web tool, which together with the report provides a user-friendly realization of complementing current annotation methods.

## 2 Related Work

Accurate genome annotation is crucial for inferring the predicted functions of an organism. Typical prokaryotic genome annotation pipelines, such as `Bakta`, `Prokka` or `PGAP`, focus on comprehensive annotation of protein-coding genes as well as house-keeping non-coding RNAs. These often lack an annotation of respective 5’ and 3’ UTR regions, as well as small ncRNAs. Other approaches, such as `Promotech` [6] or `G4promfinder` [9], complement these predictions by characterizing promoter regions. However, the diversity of regulation mechanisms makes it difficult to generalize such approaches. Adding the transcriptomic layer of information provides more specific results, for example to identify further genes or to decipher the exact transcript structure. Tools such as `APERO` [21] and `baerhunter` [24] integrate transcriptomic data with genome data by using mapped reads from RNA-seq experiments to find covered regions that are not close to annotated genes and identify putative small non-coding genes. However, RNA-seq data alone has been proven not to be a solid foundation for the base-exact prediction of gene boundaries [3]. Instead of relying only on RNA-seq data, `ANNOgesic` [34] uses dRNA-seq data to identify TSSs together with their corresponding transcripts. `ANNOgesic` offers a plethora of different methods to enhance bacterial genome annotations, including the prediction of small RNAs (sRNA), small open reading frames (sORF), circular RNAs, CRISPR related RNAs, riboswitches and RNA-thermometers. However, `ANNOgesic` does not provide a user-friendly interface both for the execution and exploration of the results to confirm and further explore the obtained predictions. Our approach, `TSS-Captur`, complements `ANNOgesic` by providing an intuitive web-based platform for exploration.

## 3 The TSS-Captur Pipeline

`TSS-Captur` is a pipeline that characterizes genomic regions starting with TSS signals that cannot be associated with any known annotated gene. It is based on `Nextflow DSL2` [10] for the stage management and `Docker` for the containerization of all required tools. To facilitate the user experience, `TSS-Captur` can be used as a web-application (see *Availability* section) and reports an interactive web-interface for result exploration (Supp. Figures S1 and S2). `NextJS` was used for client-rendering and back-end implementation, while the report is based on `Frozen-Flask` and the JavaScript library `DataTables`. Furthermore, `TSS-Captur` can also be used as a command-line pipeline.

Here, we first provide a short overview of the characterization steps (see also Fig. 1), the respective steps are then explained in detail below. Three different input files are required for `TSS-Captur`: the `MasterTable` from the TSS determination and classification process run by `TSSpredator`, as well as the genome and annotation files used for the computation of the TSSs. `TSS-Captur` first filters all TSSs classified as oTSSs or aTSSs from the `MasterTable`, extracts the respective downstream genomic and then classifies the region as either coding or non-coding. For this, two different methods are used: `CNIT` (a support-vector-based method) [14] as well as `QRNA` [25], a comparative sequence analysis requiring a pairwise alignment. Furthermore, genome-wide transcription termination sites (TTS) are predicted using the two tools `TransTermHP` [20] and `RhoTermPredict` [9]. For each predicted transcript, a putative TTS is allocated. In the case of transcripts classified as ncRNA genes, `TSS-Captur` computes their secondary structure using `RNAFold` [13]. Furthermore, the putative promoter regions of all transcripts are analyzed using `MEME` [2] to identify enriched motifs. All results are summarized in an interactive interface for easy exploration.

**Figure 1:**
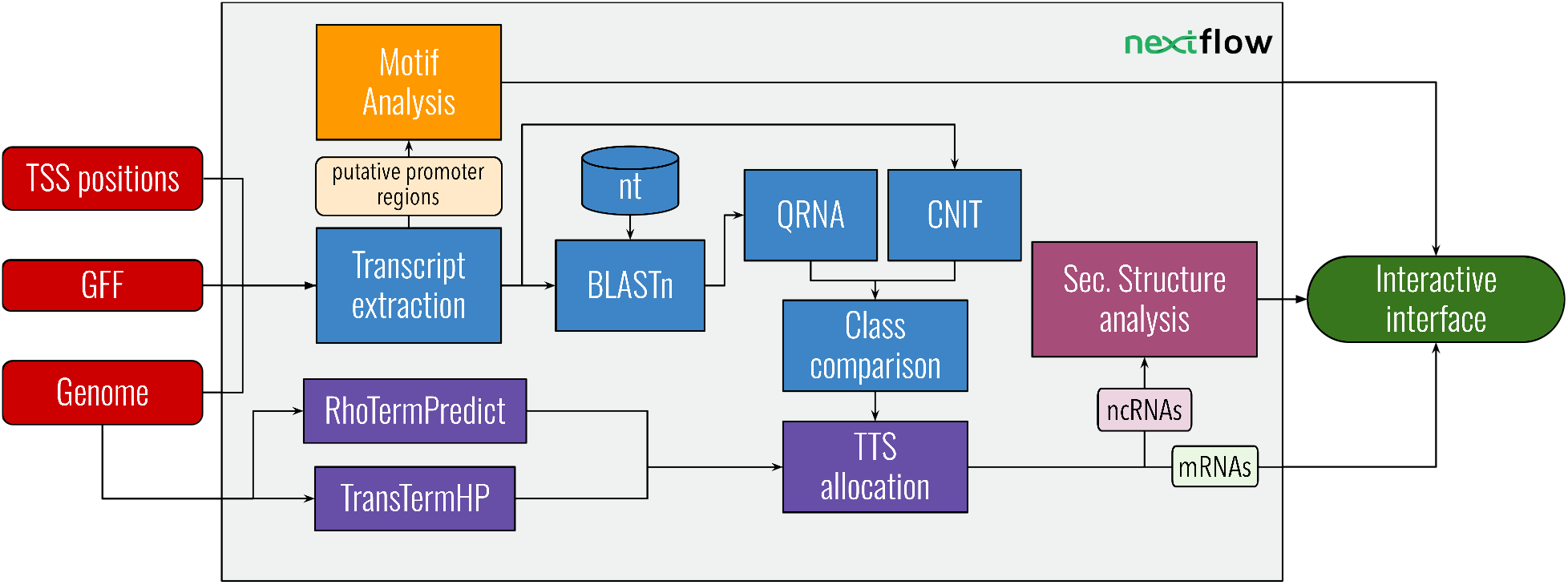
Overview of `TSS-Captur`. Different shapes have been used to represent data (ovals), processes (rectangles) or databases (cylinders). The input files (red) are provided to `Nextflow` (light green) for the data preparation. This provides the input to the RNA classification (light blue) and the motif analysis (yellow). The transcription termination site (TTS, purple) prediction works directly on the genome and the results are combined with the classified transcripts. Here, transcripts classified as ncRNAs are analyzed for their secondary structure (magenta). Lastly, all results are presented in the interactive interface (green).

### 3.1 Data Preparation

First, the `MasterTable` computed using `TSSpredator` is filtered for all TSSs classified as orphan TSSs (oTSS) or antisense TSSs (aTSS). From there, sequences downstream of these TSSs are extracted from the genome. To approximate a transcript length, the mean gene length (*µ*_*G*_) is calculated from all annotated genes. This *µ*_*G*_ serves as the transcript length’s upper limit, with a maximum of 1000bp to fulfill the limitations of `QRNA`. Transcript extraction proceeds from each TSS until this upper limit is reached, or another genomic feature is encountered. These features include any TSS present in the `MasterTable` (either primary, secondary, antisense or orphan) or the start of an annotated gene. If an extracted region is smaller than 30 bp, it is removed from the analysis together with its TSS. The lower limit of 30 bp follows the recommendations from Rivas and Eddy (2001), and models also the expected minimal length of ncRNAs [25, 30]. Following extraction, the transcripts are saved in `FASTA`-files, one file for each strand together with their TSS class (oTSS or aTSS). To ensure that all transcripts have the same reading direction, the reverse complement is taken for those extracted from the reverse strand.

### 3.2 Classification of transcripts

For the characterization of transcripts, `TSS-Captur` uses the comparative genomic tool `QRNA` and the tool `CNIT` [25, 14]. `CNIT` is an *ab initio* characterization tool based on SVMs that requires only the transcript itself as input. It provides two pre-trained models: one for animals and one for plants. To our knowledge, no similar tool has been developed for prokaryotes. We tested both models based on a subset of annotated genes (see *Verification* and *Use Case* below), and found its performance satisfactory also for prokaryotes.

While `CNIT` classifies a sequence *ab initio*, `QRNA` relies on a pairwise alignment. To determine the most appropriate alignment for each sequence, `TSS-Captur` uses `BLAST` and the `nt` database. For optimal evolutionary distance (which is recommended to be 65-85% similarity), the sensitive `dc-Megablast` search strategy of `BLAST` is employed, since it mainly returns hits with 80% similarity [23]. To address the longer runtime of this sensitive search, `TSS-Captur` reduces the species considered within the `BLAST` search to closely related ones. This is achieved by using the organism’s accession code and the `ETE toolkit` [18] library to query the `NCBI Taxonomy`. The hits reported by `BLAST` are evaluated by prioritizing that: (1) the similarity is within the optimal range and (2) the coverage of the transcript is maximized, with a special focus on the 5’-region of the query (see Supplementary Section B for details). For each transcript, the best alignment is stored in a `FASTA`-file for usage in `QRNA`.

Running both classifiers, `CNIT` and `QRNA`, allows accounting for different intrinsic characteristics of the sequence and evolutionary aspects for the classification and thus strengthens the prediction. The output of both tools are compared to compute a final call for each transcript. Both tools provide a class prediction, a prediction score, and the best functional genomic region (e.g., the CDS region for coding sequences). Hence, these tools already provide putative 3’-regions as well as possible TTS. `TSS-Captur` computes a class-specific *z*-score normalization to enable a comparison of the two methods’ prediction scores. In case of agreement, the output of the best-scoring program is used. In the case of class disagreement of the two tools, `TSS-Captur` proceeds as follows:

- If either of the tools predicts a coding gene, and the other a ncRNA, we prioritize the decision for the coding gene (see *Verification* for details).
- If `QRNA` returns no prediction due to a missing alignment or labels the transcript as *OTH* (null model), `CNIT`’s prediction is used.

The final output of the chosen program are coordinates that define the boundaries of the transcript, including a putative 3’-end. This end will be refined by identifying a nearby putative TTS in the following step.

### 3.3 Transcription termination sites

`TSS-Captur` incorporates termination site prediction in the characterization, utilizing `TransTermHP` [20] to identify intrinsic termination and `RhoTermPredict` [9] for rho-dependent termination genome-wide. `TransTermHP` is run using `nocoRNAc` [17] as a wrapper, since it is optimized for whole-genome analysis and directly filters terminators with low confidence scores (below 75). Both tools report `GFF`-files containing genome-wide predicted termination regions with a confidence score.

After transcript classification as explained above, `TSS-Captur` associates transcripts with potential terminators by comparing the coordinates as predicted by either `QRNA` or `CNIT` with the results of the termination site prediction. Each termination site is evaluated by considering its confidence score and its distance to the 3’-end predicted during classification (see Supplementary Section C for details). The best-scoring terminator is taken for each transcript. Finally, `TSS-Captur` adjusts the transcript’s TTS based on the type of terminator allocated to a transcript. For intrinsic terminators, the end of the hairpin loop marks the TTS. In rho-dependent cases, the RNAP pause site (150 nt downstream of the RUT site) is analyzed, since it can either contain a hairpin or only a RNAP-pause site [16]. If a hairpin structure is detected, both the RUT site and hairpin are included in the termination region, with the TTS placed directly behind the hairpin. If no hairpin is found, the end of the RNAP pause site is considered as the transcript’s end. This returns for each transcript a TTS and a transcript length (TTS position − TSS position). After this step, a transcript’s final length may be shorter than the 30 nt threshold used in the beginning for the transcript extraction.

### 3.4 Secondary structure prediction

After the characterization of the transcripts’ ends, those classified as ncRNA are analyzed for their secondary structure. For this, `RNAFold` [22] is used, which analyzes the different possible structures for the transcript and returns the one with the lowest minimum free energy (`MFE`). Each RNA structure is saved as a `JPG`-file and shown in the resulting report table.

### 3.5 Promoter region analyses

In addition to the characterization of the transcripts, the putative promoter region of each accounted TSS is analyzed using `MEME` [2] to identify over-represented motifs. This helps identify known transcription factor binding sites, such as the Pribnow-Box or the −35 box, and hence strengthening the validity of TSS signals, as well as other possible novel regulatory elements. To achieve this, `TSS-Captur` extracts the 50 nt upstream region of each TSS to be passed to `MEME`, which identifies a user defined number of motifs with a length between 5 and 20 nt. As long as no more than 1,000 TSSs are analyzed in one batch, `MEME` will also provide the start of each motif in each sequence, which is relevant to identify common regulation motifs. Results are provided by `MEME` in a user-friendly `HTML` report and as an `XML`-file. The `XML` output is further converted into a `TSV`-file for easier data manipulation. Both report formats are included in `TSS-Captur`’s final output.

### 3.6 TSS-Captur Result Interface

To streamline exploration of `TSS-Captur`’s output, a comprehensive `HTML` report summarizes all results in a common interface (see Supplementary Figure S2). The interface visualizes all results using an interactive table for each characterization step. The *overview* page offers a summary of skipped and analyzed TSSs, including their classification as either orphan or antisense. A central table on this page merges important data: TSS position, strand, predicted class (coding/non-coding), promoter motifs, among other information. The `HTML`-format easily allows the integration of secondary structure images for each transcript. Dedicated pages for *terminator allocation* and *classification* provide in-depth information on these processes. For example, the classification page allows users to trace the rationale behind each transcript’s assigned class. Direct integration of the `MEME HTML` report offers detailed inspection of promoter motifs, leveraging all of `MEME`’s existing visualization advantages. Finally, the *ignored TSS* page lists TSSs excluded by `TSS-Captur` due to their short length.

## 4 Verification of TSS-Captur

To assess the overall performance of `TSS-Captur`, we used a well-known and established dataset and its corresponding annotations from *Campylobacter jejuni* str. NCTC 11169 (RefSeq ID NC_002163). The `MasterTable` was computed using the data published by Dugar *et al*. [11] and `TSSpredator` with default parameters. We selected a subset of 100 primary TSSs (pTSS) and their corresponding annotations, equally balancing both RNA classes (ncRNA and mRNAs).

First, we extracted the transcripts using the exact coordinates of each annotated gene of the subset to assess the two models (animal-based and plant-based) of `CNIT` with respect to sensitivity, specificity and precision. We found that for all three metrics, the *plant* model delivered the best results (at least 0.92, see Table 1),

**Table 1:** Evaluation of CNIT’s performance on *C. jejuni*’s subset. Each model was evaluated by computing the sensitivity, specificity and precision concerning the real RNA class.

| Model | Type | Sen. | Spec. | Prec. |
| --- | --- | --- | --- | --- |
| Plant | COD | 0.92 | 1 | 1 |
|  | RNA | 1 | 0.92 | 0.926 |
| Animal | COD | 0.82 | 1 | 1 |
|  | RNA | 1 | 0.82 | 0.847 |

Afterwards, the influence of the extracted transcript’s length on the overall RNA classification implemented in `TSS-Captur` was assessed. For this, for each TSS and its annotated transcript end *l*, multiple regions with varying end *l*^′^ were extracted, with *l*^′^ = *l* + *t* and *t* = 5*i, i* ∈ [−20, 20]. To achieve a larger dataset, we lowered the 30 nt extraction threshold to 5 nt for this analysis. Based on the RNA class (mRNA or ncRNA) from the annotation file, we computed the precision for the three classification approaches (`CNIT`, `QRNA` and `TSS-Captur`’s resulting decision of the combination of both methods). This analysis revealed that the combination of both approaches implemented in `TSS-Captur` had the highest precision across most of the transcripts for ncRNA genes, while `CNIT`’s precision was the highest for mRNA genes across all lengths (Fig. 2). Still, our implementation returned precise results (*>* 0.95) across all transcripts’ length.

**Figure 2:**
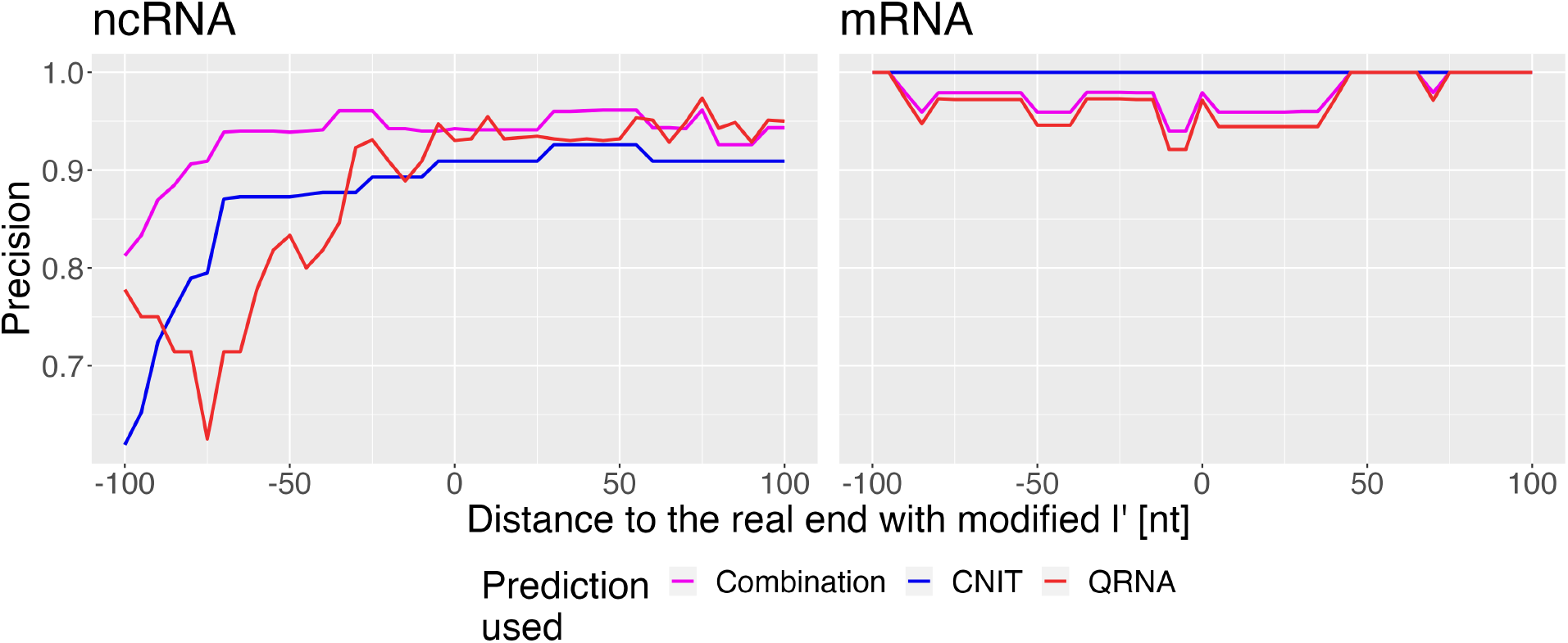
Evaluation of RNA classification. The three different types of predictions are shown: `CNIT`, `QRNA` and the combination implemented in `TSS-Captur`. For each prediction type and for each modified *l*^*′*^, the precision was computed for each type of transcript (ncRNA and mRNA).

Lastly, we modified the `MasterTable` to assess the whole process behind `TSS-Captur`. We first removed all oTSSs or aTSSs from the `MasterTable` to avoid their analysis in `TSS-Captur`. The entries in the `MasterTable` and the annotation file for the 100 pTSSs of the subset were modified to simulate orphan TSSs. The remaining TSSs were left unmodified to have a feasible `MasterTable`. The output of `TSS-Captur` was compared to the annotated classes and length for evaluation. All ncRNA genes were correctly classified, and of the 50 protein-coding genes, only three were classified as ncRNAs (Fig. 3A). For the prediction of the gene’s length, the difference between the predicted and the annotated length was computed, returning a mean difference of 9.2 nt and a median of −35 nt (Fig. 3B). 57 genes had an absolute length difference ≤ 100 nt, while 74 genes had a length difference ≤ 250 nt. For those 18 genes with an absolute length difference ≥ 500 nt, a clear difference in the classes was observed. On the one hand, eight coding genes were strongly underestimated in their length. On the other hand, 10 ncRNAs were greatly overestimated in their length.

**Figure 3:**
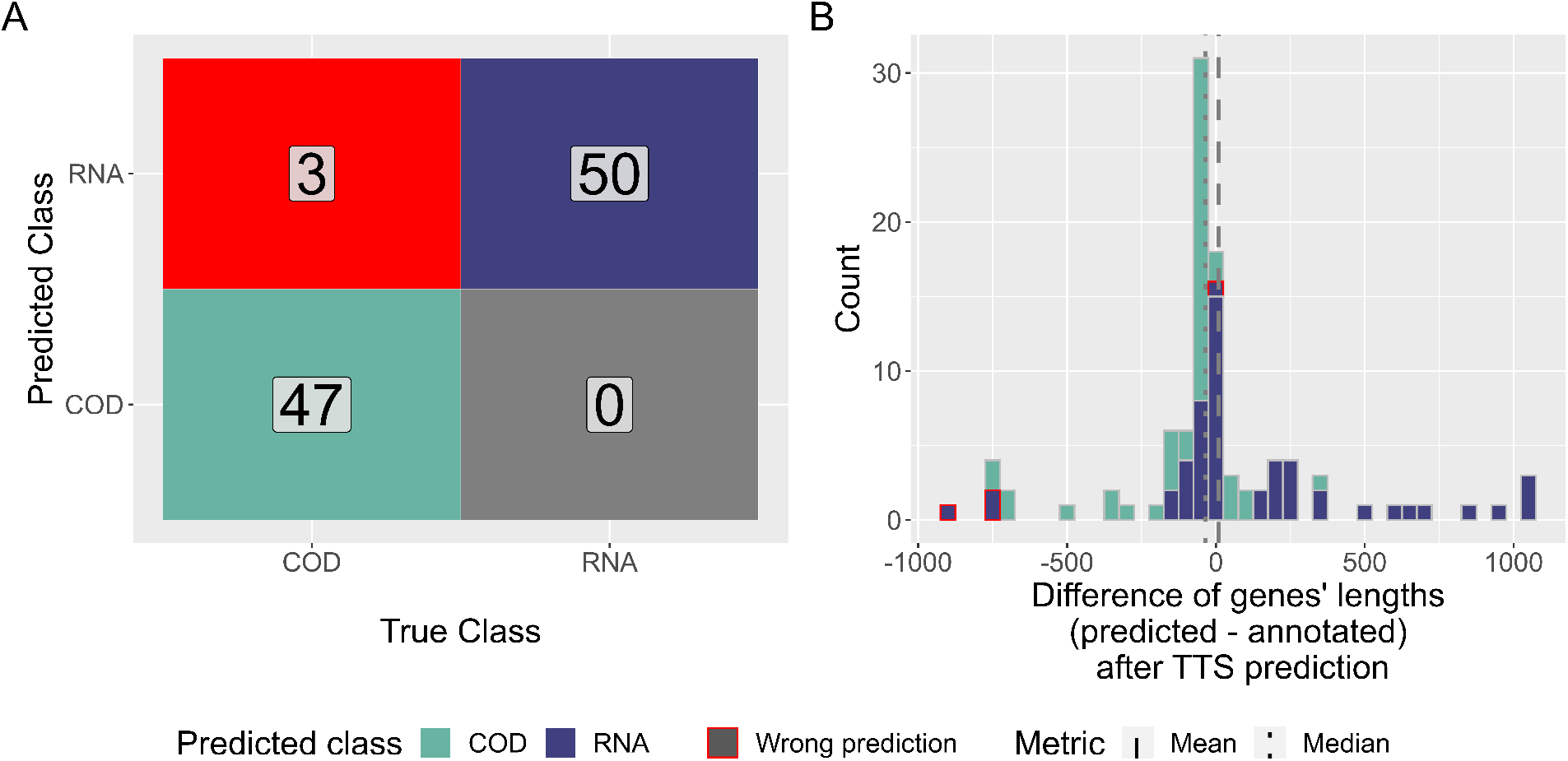
Evaluation of `TSS-Captur` by treating pTSSs as oTSSs. (A): Confusion matrix on the real and predicted RNA classes for transcript. (B): Distributions of the genes’ differences (predicted - annotated) by predicted RNA class (bin width = 50 nt). The mean and median differences are also shown. Wrongly predicted RNAs are highlighted (red).

## 5 Use Case: *Streptomyces coelicolor*

To demonstrate the applicability of `TSS-Captur`, we analyzed a genome-wide TSS dataset of *Streptomyces coelicolor* [19]. For this, `TSSpredator` was run with sensitive parameters, resulting in a total of 1, 660 antisense and orphan TSSs that were passed to `TSS-Captur` as input.

118 of these TSSs were filtered (i.e. too short transcripts), of the remaining 1542 TSSs, 927 regions were predicted as ncRNA genes and 615 as coding genes (Fig. 4A, Supplementary Fig. S2). The majority of the analysed transcripts were predicted to have a rho-dependent terminator site (80.9%), 2.7% showed intrinsic termination and for 14.4% of the sequences no terminators were predicted (Fig. 4B). The predicted ncRNA transcripts as well as coding genes displayed a wide range of lengths (Fig. 4D). With a median length of 940 nt, coding genes are on average around twice as long as ncRNAs, which show a median length of 495 nt. The secondary structure of genes predicted as ncRNAs was also computed and included in the report (see Supplementary Fig. S3).

**Figure 4:**
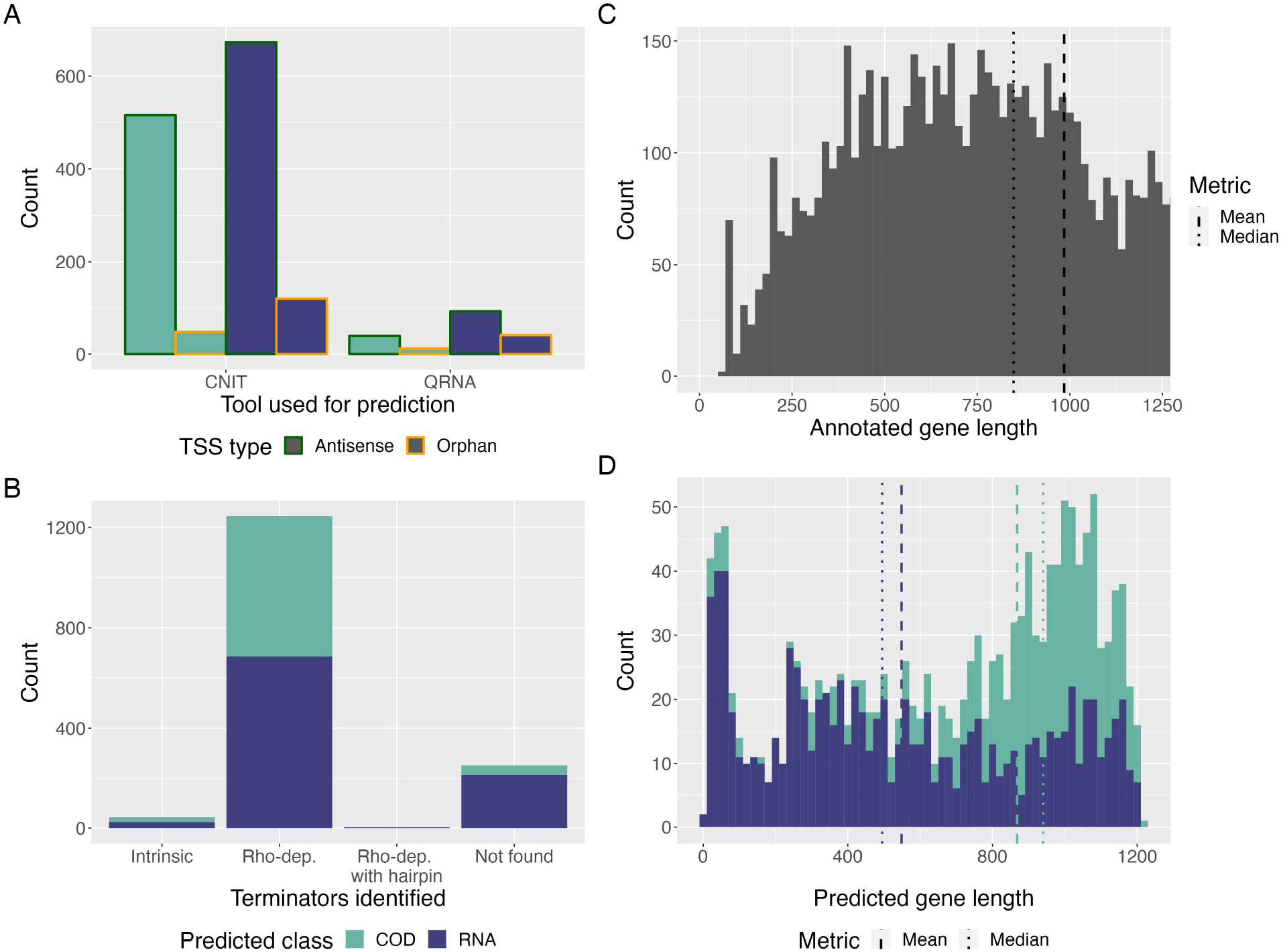
Visual summary of `TSS-Captur` output for *S. coelicolor*’s data. (A): Summary of the predicted RNA classes for each transcript by the tool used for prediction and the original TSS type. (B): Distribution of the allocated terminator regions, wrt. their type. (C & D): Histogram (bin width = 20) of the length for annotated genes (C) and for predicted transcripts (D). Only annotated genes shorter than 1250 nt are shown for a better comparison with the predictions.

`TSS-Captur` identified two enriched motifs within the upstream regions of the analyzed transcripts. The most-significant motif (*E*-value = 1.9 × 10^−438^) was found in 589 promoter sequences, exhibiting a strong G/C pattern and a preference for position 20 nucleotides upstream of the TSS (Fig. 5). A database comparison using `TOMTOM` [15] was started from the report of `TSS-Captur`. However, it did not return any matches for this motif.

**Figure 5:**
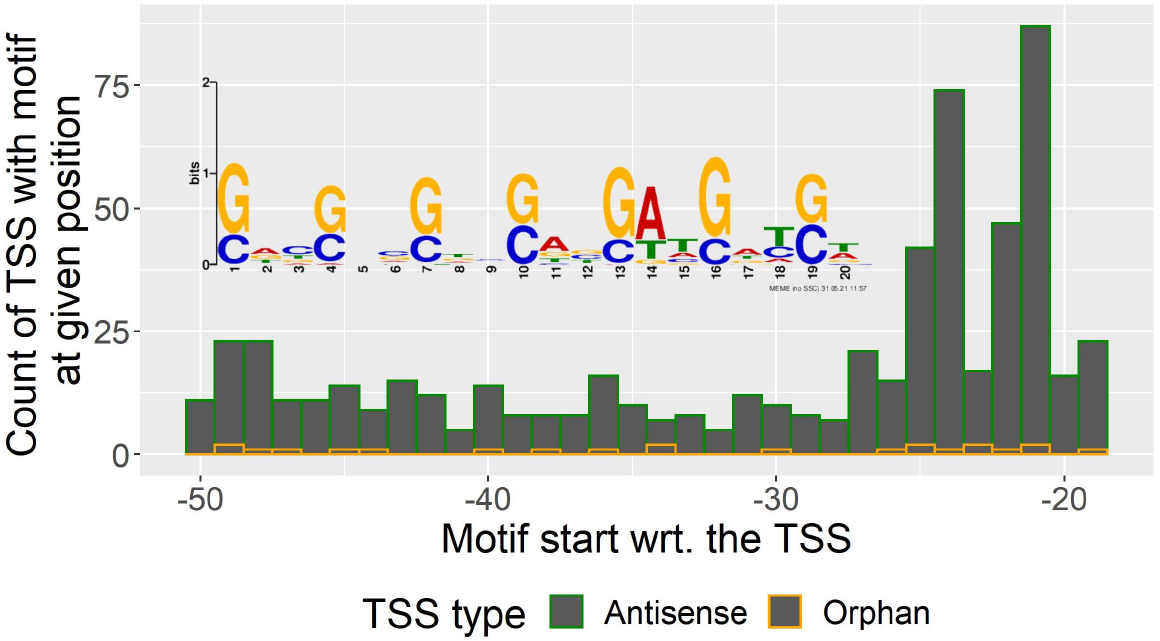
Distribution of start of most enriched motif in the promoter regions for *S. coelicolor*’s analyzed TSSs using `MEME`.

Another study by Vockenhuber *et al*. [33] identified 63 putative small ncRNA genes based on RNA-seq and applied Northern Blot to eleven of them, confirming this subset. We compared the 63 putative ncRNAs as predicted by Vockenhuber *et al*. to the results generated by `TSS-Captur` based on the study by Jeong *et al*. We reduced the sets of TSSs computed with Jeong *et al*.’s data to those that had a distance of *±*15 nt to the regions reported by Vockenhuber *et al*. This comparison revealed 23 intersecting TSS positions, including six that were confirmed using Northern Blot (Supplementary Table S1). Of the other 40 TSSs, 10 were classified as primary TSSs by `TSSpredator`, while 30 were not listed in the reduced subset. Of the overlapping 23 transcripts, `TSS-Captur` classified 18 as ncRNA genes, including the six genes confirmed using Northern Blot. Compared to the lengths reported by the study, `TSS-Captur` predicted longer transcripts (Supplementary Fig. S4). To further validate our predictions, we compared all transcripts classified as ncRNA genes against the `RFAM` database. This returned 11 unique hits (Supplementary Table S2), of which three hits confirmed transcripts identified also in the study by Vockenhuber *et al*.

## 6 Runtime analysis

The runtime of `TSS-Captur` was assessed using multiple datasets (in total, 18 different runs were conducted) and includes all genome-wide analyses such as the termination site predictions and promoter analyses. It was performed on a server with an Intel(R) Xeon(R) E5-4610 v2 @ 2.30GHz CPU using at most 4 cores. The results show a clear linear dependency on the number of TSSs (Figure 6, left) with an increase in the runtime by 18.6s per TSS. In all runs, the most time-consuming step of 89.5% is needed by `QRNA` (see Figure 6, right).

**Figure 6:**
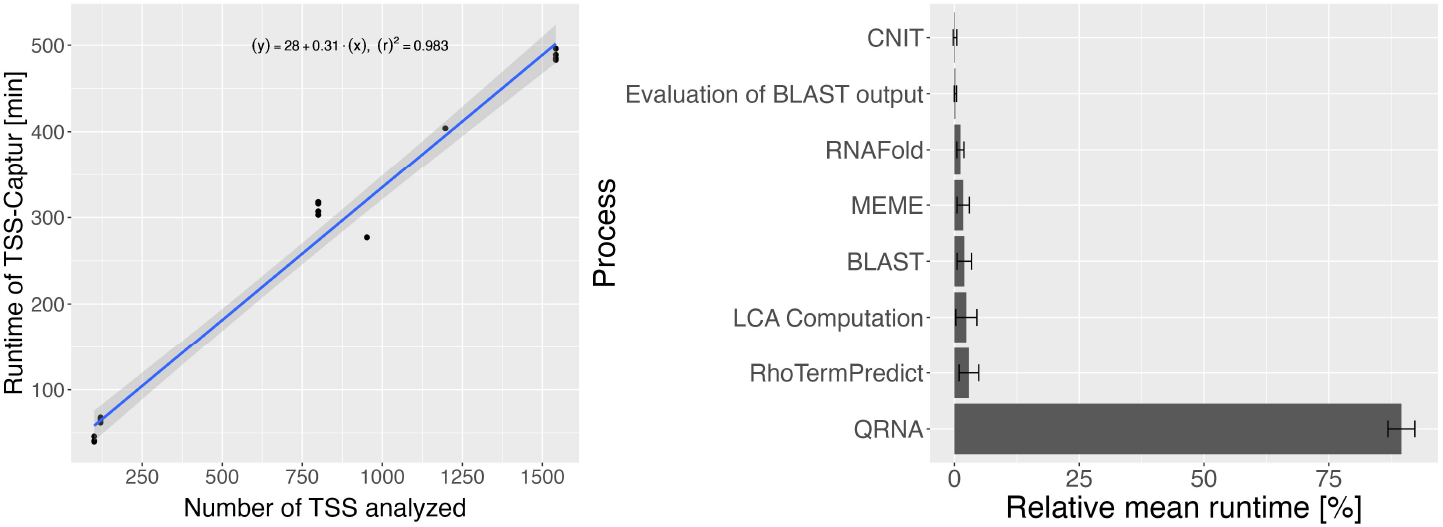
Runtime analysis of `TSS-Captur`. The runtime of 18 analyses is visualized together with the computed linear regression line (left). The mean of the runtime as well as standard deviation of each process was computed and shown here in % relative to the overall runtime (right). Processes that were faster than one minute are not shown.

## 7 Discussion and Outlook

Genome-wide base-resolved transcriptome maps offer insights into regulation regions of annotated genes (e.g., 5’-UTRs, promoters) [3, 26]. However, they also reveal strong transcription signals in unannotated regions – a rich source of potentially undiscovered genes. Our exploration tool, `TSS-Captur`, unlocks the potential of these sites, enabling user-friendly analysis that combines genomic and transcriptomic data for the characterization of these sites.

The main back-end component of `TSS-Captur` is a pipeline implemented using `Nextflow`. Among the advantages of `Nextflow` are its declarative syntax, as well as modularity and extensibility. Moreover, our approach covers multiple aspects of the characterization. For the classification of uncharacterized transcript sequences, both intrinsic genomic features (`CNIT`) and evolutionary conservation (`QRNA`) are considered. Similarly, the transcription termination site prediction combines complementary methods (`TransTermHP & RhoTermPredict`) for a broad analysis. The analysis of secondary structures and promoter regions offers insights into the function of the transcript as well as its regulation mechanisms. A unified report facilitates exploration of the results.

To validate our approach, we used a dataset based on primary TSSs in *Campylobacter jejuni*. We analyzed the precision of the RNA classification across varying transcript lengths, demonstrating that our combined approach provides a balanced performance for both gene classes. By simulating orphan TSSs, we evaluated the overall performance of `TSS-Captur`. The results showed accurate classification for all ncRNAs and almost all protein-coding genes, while lengths’ predictions vary in accuracy across genes.

Our analysis of the *S. coelicolor* TSS dataset showcases the utility of `TSS-Captur` in exploring unannotated genomic regions. Of particular interest is the identification of a highly enriched G/C motif upstream of many of the unannotated transcripts. While no direct matches were found using `TOMTOM`, such a motif could be a signal of a G-quadruplex motif that has been linked to regulation mechanisms in *S. coelicolor* [8]. The user could also start a motif-search using `FIMO` [1] from the report of `TSS-Captur` to search for this motif in all promoter regions. This finding demonstrates how `TSS-Captur` facilitates hypothesis generation and links to external tools for deeper analysis.

`TSS-Captur`’s predictions exhibited some overlap with the experimentally validated ncRNAs from Vockenhuber *et al*. [33]. Here, it is important to remark that the experimental conditions between the analyzed TSSs with `TSS-Captur` and the study of Vockenhuber *et al*. differed, hence not all transcripts might be present. This, together with our choice of using the TSS prediction of TSSpredator with the most sensitive parameters, explains the rather low rate of commonly called TSSs. Nonetheless, the confirmation with the study, along with the `RFAM` validation of some predicted ncRNAs, emphasizes how `TSS-Captur` could be used to detect novel ncRNAs.

When comparing `TSS-Captur` and `ANNOgesic`, both tools use a number of different tools for the various aspects of transcript annotation in their backends. While `ANNOgesic` currently offers a larger range of methods than `TSS-Captur`, addressing additional aspects of transcriptomic data annotation, it also has some limitations. `ANNOgesic`’s reliance on existing databases and known mechanisms (like the Shine-Dalgarno sequence) for transcript characterization may limit the discovery of novel genes. In contrast, `TSS-Captur`’s classification relies on more abstract characteristics of the transcripts, with the potential of uncovering unknown mechanisms. Moreover, `ANNOgesic`’s command-line interface poses a usability challenge for users in the field who might not have extensive computational skills, especially due to the long documentation required to understand the program. One motivation to develop `TSS-Captur` was to provide users with a user-friendly and accessible tool. Our web-interface simplifies file parsing, eliminating the need for direct command-line interaction for inexperienced users. Moreover, `TSS-Captur` unifies results within a single interface, offering data exploration alongside convenient access to secondary structure visualizations and external analysis options, such as `MEME`.

Due to the flexibility of `Nextflow`, new modules, tools or even other data sources can be easily included in `TSS-Captur`. One issue that we identified in our two use cases is the prediction of the 3’-end. In particular, the lengths of predicted ncRNAs seem often overestimated. This was revealed by the validation study as well as our observation that `TSS-Captur`’s predicted transcripts often aligned only with subsequences within the `RFAM` entries, indicating an overestimation of the predicted length. In the future, we may consider integrating more recent tools or other data sources for the prediction of the 3’-end, such as `TermNN` [4] or Term-seq data [7]. For the classification step of uncharacterized transcript regions, we used two complementary methods. However, here we still see an opportunity for further developments of classifying transcripts in prokaryotes. Such new tools can then be easily integrated in `TSS-Captur`.

In summary, starting from experimental TSS data, `TSS-Captur` predicts the characterization of unclassified signals and allows a user-friendly exploration to complement prokaryotic annotation tools, contributing to the understanding of bacterial transcriptomes. By combining a user-centric design, `TSS-Captur` empowers researchers with diverse technical backgrounds to characterize experimentally detected transcription signals and eventually accelerate the discovery of novel genomic elements and their regulatory mechanisms.

## 8 Availability

`TSS-Captur` can be accessed at https://tsscaptur-tuevis.cs.uni-tuebingen.de/. The source code is available via `Zenodo` at https://zenodo.org/doi/10.5281/zenodo.12527008 and via `GitHub` athttps://github.com/Integrative-Transcriptomics/tss-captur-dsl2. The datasets used for this study are also available online at https://zenodo.org/doi/10.5281/zenodo.12526907. All tools required for the pipeline have been containerized into a Docker image and are publicly available at https://hub.docker.com/r/mwittep/tsscaptur

## 9 Competing interests

No competing interest is declared.

## 10 Author contributions statement

MW and KN conceived the idea for `TSS-Captur`. MW implemented the first pipeline version and conducted the experiments. TV implemented the `DSL2` version and the web application. All authors wrote and reviewed the manuscript.

## 11 Funding

We acknowledge infrastructural funding from the Cluster of Excellence EXC 2124 ‘Controlling Microbes to Fight Infections’ [project ID 390838134] from the German Research Foundation (DFG).

## Notes

https://zenodo.org/doi/10.5281/zenodo.12527008

https://zenodo.org/doi/10.5281/zenodo.12526907

https://tsscaptur-tuevis.cs.uni-tuebingen.de/

### Competing Interest Statement

The authors have declared no competing interest.

## References

[1] T. L. Bailey, M. Boden, F. A. Buske, M. Frith, C. E. Grant, L. Clementi, J. Ren, W. W. Li, and W. S. Noble. MEME Suite: tools for motif discovery and searching. Nucleic Acids Research, 37(suppl_2):W202–W208, 2009.

[2] T. L. Bailey and C. Elkan. Fitting a mixture model by expectation maximization to discover motifs in biopolymers. Proceedings. International Conference on Intelligent Systems for Molecular Biology, 2:28–36, 1994.

[3] T. Bischler, H. S. Tan, K. Nieselt, and C. M. Sharma. Differential RNA-seq (dRNA-seq) for annotation of transcriptional start sites and small RNAs in Helicobacter pylori. Methods, 86:89–101, 2015.

[4] V. B. Brandenburg, F. Narberhaus, and A. Mosig. Inverse folding based pre-training for the reliable identification of intrinsic transcription terminators. PLOS Computational Biology, 18(7):e1010240, July 2022.

[5] C. Carvalho Barbosa, S. H. Calhoun, and H.-J. Wieden. Non-coding rnas: what are we missing? Biochemistry and Cell Biology, 98(1):23–30, 2020.

[6] R. Chevez-Guardado and L. Peña-Castillo. Promotech: a general tool for bacterial promoter recognition. Genome Biology, 22(1):318, Nov. 2021.

[7] D. Dar, M. Shamir, J. R. Mellin, M. Koutero, N. Stern-Ginossar, P. Cossart, and R. Sorek. Term-seq reveals abundant ribo-regulation of antibiotics resistance in bacteria. Science, 352(6282):aad9822, 2016.

[8] M. Di Salvo, E. Pinatel, A. Talà, M. Fondi, C. Peano, and P. Alifano. G4promfinder: an algorithm for predicting transcription promoters in gc-rich bacterial genomes based on at-rich elements and g-quadruplex motifs. BMC Bioinformatics, 19(1):36, Feb. 2018.

[9] M. Di Salvo, S. Puccio, C. Peano, S. Lacour, and P. Alifano. RhoTermPredict: an algorithm for predicting rho-dependent transcription terminators based on Escherichia coli, Bacillus subtilis and Salmonella entericadatabases. BMC Bioinformatics, 20(1):117, 2019.

[10] P. Di Tommaso, M. Chatzou, E. W. Floden, P. P. Barja, E. Palumbo, and C. Notredame. Nextflow enables reproducible computational workflows. Nature Biotechnology, 35(4):316–319, 2017.

[11] G. Dugar, A. Herbig, K. U. Förstner, N. Heidrich, R. Reinhardt, K. Nieselt, and C. M. Sharma. High-resolution transcriptome maps reveal strain-specific regulatory features of multiple Campylobacter jejuni isolates. PLOS Genetics, 9(5):e1003495, 2013.

[12] L. Ettwiller, J. Buswell, E. Yigit, and I. Schildkraut. A novel enrichment strategy reveals unprecedented number of novel transcription start sites at single base resolution in a model prokaryote and the gut microbiome. BMC Genomics, 17(1):199, 2016.

[13] A. R. Gruber, R. Lorenz, S. H. Bernhart, R. Neuböck, and I. L. Hofacker. The Vienna RNA websuite. Nucleic Acids Research, 36(uppl_2):W70–W74, 2008.

[14] J.-C. Guo, S.-S. Fang, Y. Wu, J.-H. Zhang, Y. Chen, J. Liu, B. Wu, J.-R. Wu, E.-M. Li, L.-Y. Xu, L. Sun, and Y. Zhao. CNIT: a fast and accurate web tool for identifying protein-coding and long non-coding transcripts based on intrinsic sequence composition. Nucleic Acids Research, 47(W1):W516–W522, 2019.

[15] S. Gupta, J. A. Stamatoyannopoulos, T. L. Bailey, and W. S. Noble. Quantifying similarity between motifs. Genome Biology, 8(2):R24, Feb. 2007.

[16] T. M. Henkin. Control of transcription termination in prokaryotes. Annual review of genetics, 30(1):35–57, 1996.

[17] A. Herbig and K. Nieselt. nocoRNAc: Characterization of non-coding RNAs in prokaryotes. BMC Bioinformatics, 12(1):40, 2011.

[18] J. Huerta-Cepas, F. Serra, and P. Bork. ETE 3: Reconstruction, analysis, and visualization of phylogenomic data. Molecular Biology and Evolution, 33(6):1635–1638, 2016.

[19] Y. Jeong, J.-N. Kim, M. W. Kim, G. Bucca, S. Cho, Y. J. Yoon, B.-G. Kim, J.-H. Roe, S. C. Kim, C. P. Smith, and B.-K. Cho. The dynamic transcriptional and translational landscape of the model antibiotic producer Streptomyces coelicolor A3(2). Nature Communications, 7(1):11605, 2016.

[20] C. L. Kingsford, K. Ayanbule, and S. L. Salzberg. Rapid, accurate, computational discovery of rho-independent transcription terminators illuminates their relationship to DNA uptake. Genome Biology, 8(2):R22, 2007.

[21] S. Leonard, S. Meyer, S. Lacour, W. Nasser, F. Hommais, and S. Reverchon. APERO: a genome-wide approach for identifying bacterial small RNAs from RNA-Seq data. Nucleic Acids Research, 47(15):e88, Sept. 2019.

[22] R. Lorenz, S. H. Bernhart, C. Höner zu Siederdissen, H. Tafer, C. Flamm, P. F. Stadler, and I. L. Hofacker. ViennaRNA package 2.0. Algorithms for Molecular Biology, 6(1):26, 2011.

[23] S. McGinnis and T. L. Madden. BLAST: at the core of a powerful and diverse set of sequence analysis tools. Nucleic Acids Research, 32(uppl_2):W20–W25, 2004.

[24] A. Ozuna, D. Liberto, R. M. Joyce, K. B. Arnvig, and I. Nobeli. baerhunter: an R package for the discovery and analysis of expressed non-coding regions in bacterial RNA-seq data. Bioinformatics, 36(3):966–969, Feb. 2020.

[25] E. Rivas and S. R. Eddy. Noncoding RNA gene detection using comparative sequence analysis. BMC Bioinformatics, 2(1):8, 2001.

[26] D. Ryan, L. Jenniches, S. Reichardt, L. Barquist, and A. J. Westermann. A high-resolution transcriptome map identifies small rna regulation of metabolism in the gut microbe bacteroides thetaiotaomicron. Nature Communications, 11(1):3557, July 2020.

[27] O. Schwengers, L. Jelonek, M. A. Dieckmann, S. Beyvers, J. Blom, and A. Goesmann. Bakta: rapid and standardized annotation of bacterial genomes via alignment-free sequence identification. Microbial Genomics, 7(11):000685, 2021.

[28] T. Seemann. Prokka: rapid prokaryotic genome annotation. Bioinformatics, 30(14):2068–2069, 2014.

[29] C. M. Sharma, S. Hoffmann, F. Darfeuille, J. Reignier, S. Findeiß, A. Sittka, S. Chabas, K. Reiche, J. Hackermüller, R. Reinhardt, P. F. Stadler, and J. Vogel. The primary transcriptome of the major human pathogen Helicobacter pylori. Nature, 464(7286):250–255, 2010.

[30] R. Sorek and P. Cossart. Prokaryotic transcriptomics: a new view on regulation, physiology and pathogenicity. Nature Reviews Genetics, 11(1):9–16, 2010.

[31] D. Stazic and B. Voß. The complexity of bacterial transcriptomes. Journal of biotechnology, 232:69–78, 2016.

[32] T. Tatusova, M. DiCuccio, A. Badretdin, V. Chetvernin, E. P. Nawrocki, L. Zaslavsky, A. Lomsadze, K. D. Pruitt, M. Borodovsky, and J. Ostell. Ncbi prokaryotic genome annotation pipeline. Nucleic Acids Research, 44(14):6614–6624, Aug. 2016.

[33] M.-P. Vockenhuber, C. M. Sharma, M. G. Statt, D. Schmidt, Z. Xu, S. Dietrich, H. Liesegang, D. H. Mathews, and B. Suess. Deep sequencing-based identification of small non-coding RNAs in Streptomyces coelicolor. RNA biology, 8(3):468–477, 2011.

[34] S.-H. Yu, J. Vogel, and K. U. Förstner. ANNOgesic: a Swiss army knife for the RNA-seq based annotation of bacterial/archaeal genomes. GigaScience, 7(9), 2018.

