## Supplementary Figures and Tables for "TSS-Captur: A User-Friendly Characterization Pipeline for Transcribed but Unclassified RNA transcripts"

#### A The TSS-Captur pipeline

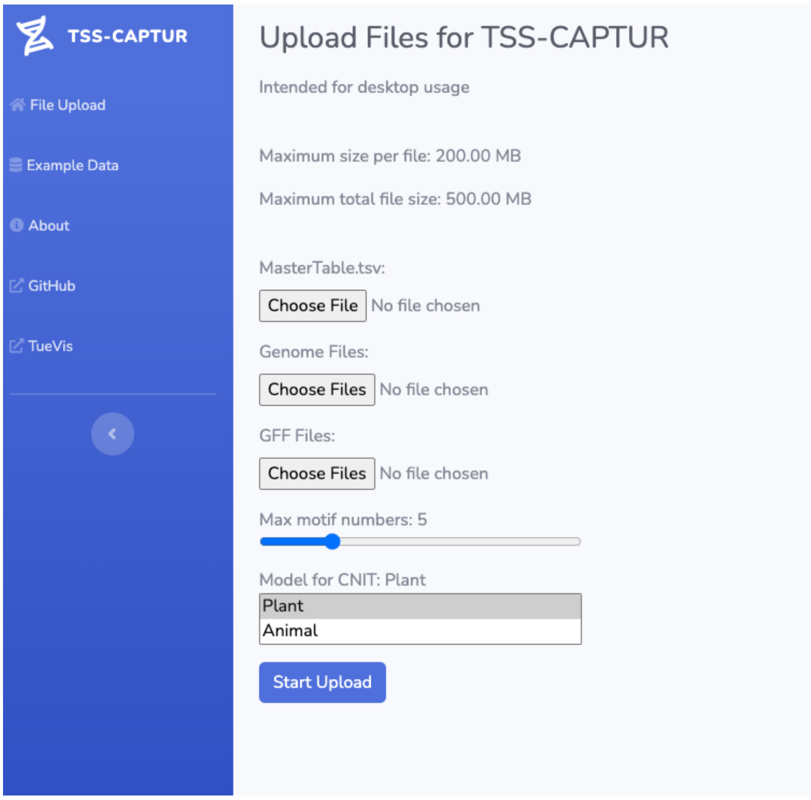


Figure S1: Snapshot of the web-interface of TSS-Captur. It allows the user to upload their files, as well as selecting a few parameters for the pipeline. Two other pages provide more information wrt. TSS-Captur. The *Example Data* page links different datasets that can be used directly in TSS-Captur, as well as their precomputed reports. The *About* page provides an overview of the pipeline, such as the tools used for characterization.

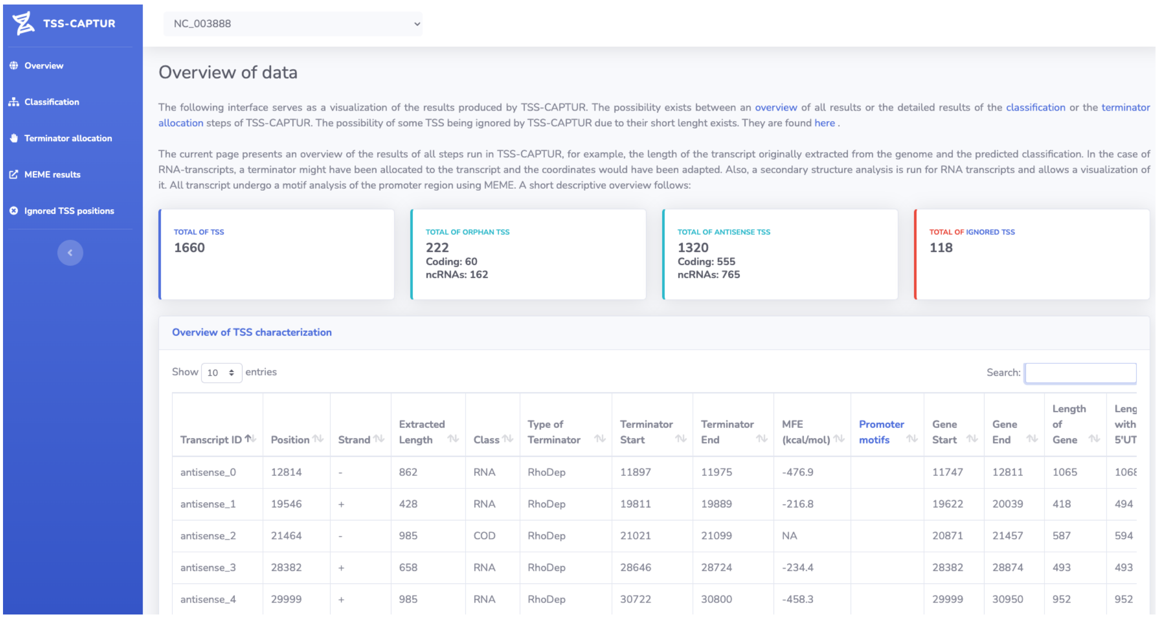


Figure S2: Snapshot of the interactive report of `TSS-Captur` visualizing *S. coelicolor*'s data. The interface provides different levels of details for each analysis step run in `TSS-Captur` that can be accessed via the navigation menu (left). An overview of the data is provided by summarizing main results of the analyzed TSSs. The information regarding each transcript is listed in a tabular view that allows sorting and filtering.

#### B Evaluation of BLAST's hits

After BLAST returns a set of local alignments $A$ for each transcript, the optimal hit $H^*$ must be selected. Instead of relying solely on the best $E$-value or bit-score, other characteristics are crucial. The ideal pairwise-alignment hit should fulfill the following criteria:

- The distance to the TSS ($D_{TSS}$) should be minimal.
- The similarity $Sim$ should be within the range of $65 - 85\%$.
- The coverage of the query ($Cov_q$) in the alignment should be maximized.

To evaluate each hit ($Hi \in A$), the scoring function Eq. 1 is used. `qcovhsp`, `qstart` and `pident` are values returned by BLAST that represent the coverage of the query, the start of coordinate of the alignment in the query and the overall identity of the alignment, respectively. A weighted average $S$ combines the three scores with weights 0.45, 0.35, and 0.2, reflecting the relative importance of TSS proximity, similarity, and query coverage. For each transcript, the alignment $H^*$ is taken, with $H^* = \arg\max S(H_i), \forall H_i \in A$, and stored in a FASTA-file for usage in QRNA.

$$\begin{aligned} D_{TSS}(S_q, t) &= \frac{\min(\max(0, S_q - t), t + 20)}{t + 20} \\ D_{Sim}(I_A) &= \frac{\mathcal{N}_{\mu=75,\sigma=8}(I_A)}{\mathcal{N}_{\mu=75,\sigma=8}(75)} \\ Cov_q(C_q) &= \min(1, \frac{C_q - 50}{25}) \\ S(H_i) &= 0.45 \cdot D_{TSS}(\texttt{qstart}_{H_i}, 10) \\ &+ 0.35 \cdot D_{Sim}(\texttt{pident}_{H_i}) \\ &+ 0.2 \cdot Cov_q(\texttt{qhspcov}_{H_i}) \end{aligned} \tag{Eq. 1}$$

### C Evaluation of termination sites

Once the RNA class is predicted for each transcript, `TSS-Captur` continues with the allocation of a predicted termination site. For this, the termination regions identified by `RhoTermPredict` and `TransTermHP` that are start within the range predicted by `CNIT` and `QRNA` are considered as putative termination sites. Since the used tools might return false-positive, such as substructures of the secondary structure of an ncRNA, the predictions need to be scored to identify the most-plausible termination site. We consider two aspects for this evaluation: the confidence value $TTS_{score}$ returned by the corresponding prediction tool, and the location of the termination region wrt. the coordinates reported by the classification step ($T_{start}$ and $T_{end}$, respectively, for the start and end of the predicted transcript). Note that $T_{start}$ is the same as the TSS position. The scores of `TransTermHP` and `RhoTermPredict` have different ranges, hence they are normalized, computing $TTS'_{score}$ A weighted average is computed using the normalized score and the distances to the TSS and the $3'$-end of the transcript, as seen in Eq. 2.

$$
\begin{aligned}
D_{start} &= \frac{|TTS_{start} - T_{start}|}{length(T)} \\
D_{end} &= \frac{|TTS_{start} - T_{end}|}{length(T)} \\
score &= \frac{TTS'_{score}}{2} + \frac{D_{start}}{4} + \left(\frac{1 - D_{end}}{4}\right)
\end{aligned}
\qquad \text{(Eq. 2)}
$$

### D Use case: *S. coelicolor*

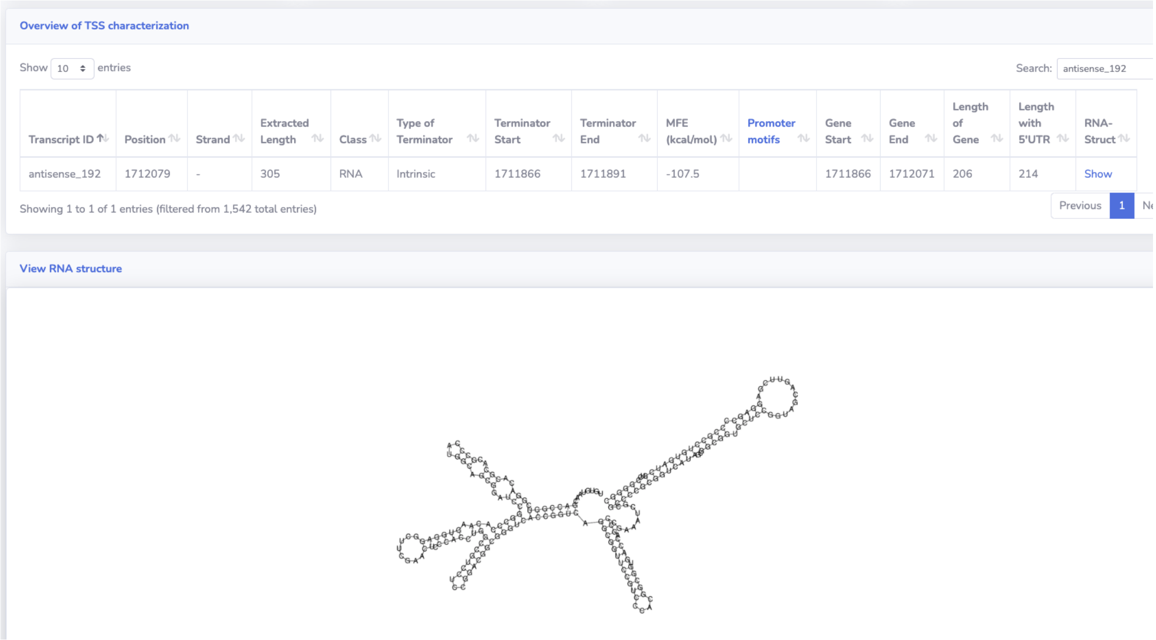


Figure S3: Interactive report of `TSS-Captur` visualizing *S. coelicolor*'s transcript *antisense_192* predicted data. The predicted secondary structure can be directly visualized in the interface by clicking the corresponding element in the table.

Table S1: Comparison of results of `TSS-Captur` with the study by Vockenhuber *et al.*. The first columns represent the name, start, end and length reported by the study. Transcripts confirmed via Northern Blot (NB) are also distinguished. For `TSS-Captur`'s results, the position of the TSS, the start, end, class and length reported by the characterization are shown.

| **Study Vockenhuber et al.** | | | | | **Results of TSS-Captur** | | | |
|---|---|---|---|---|---|---|---|---|
| **Name** | **Start** | **End** | **Length** | **NB** | **ID** | **TSS** | **Length** | **Class** |
| scr0991 | 1045891 | 1046090 | 200 | Yes | orphan_20 | 1045891 | 351 | RNA |
| scr1601 | 1711967 | 1712074 | 214 | Yes | antisense_192 | 1712079 | 214 | RNA |
| scr1821 | 1950867 | 1950948 | 82 | Yes | orphan_32 | 1950867 | 1077 | RNA |
| scr2736-1 | 2982224 | 2982505 | 282 | Yes | orphan_48 | 2982224 | 34 | RNA |
| as2780 | 3034099 | 3034209 | 111 | Yes | antisense_376 | 3034209 | 1142 | RNA |
| scr3035 | 3321216 | 3321353 | 138 | Yes | orphan_59 | 3321353 | 140 | RNA |
| as3287 | 3636488 | 3636569 | 82 | No | antisense_488 | 3636567 | 527 | RNA |
| as3317 | 3669433 | 3669657 | 225 | No | antisense_491 | 3669433 | 1104 | COD |
| as3321 | 3673385 | 3673581 | 197 | No | antisense_494 | 3673385 | 709 | COD |
| scr3323 | 3675098 | 3675329 | 232 | No | orphan_77 | 3675098 | 331 | RNA |
| scr3558 | 3933527 | 3933668 | 142 | No | antisense_599 | 3933677 | 32 | RNA |
| scr3559 | 3934693 | 3934927 | 235 | No | orphan_88 | 3934687 | 71 | RNA |
| scr3920 | 4315358 | 4315484 | 127 | No | antisense_682 | 4315355 | 848 | COD |
| scr3928 | 4323842 | 4324023 | 182 | No | orphan_109 | 4323842 | 584 | RNA |
| as4261 | 4674583 | 4674770 | 188 | No | antisense_744 | 4674582 | 36 | RNA |
| as4567 | 4984635 | 4984793 | 159 | No | antisense_804 | 4984793 | 1059 | COD |
| scr4632 | 5055055 | 5055172 | 118 | No | orphan_142 | 5055173 | 533 | RNA |
| as4675 | 5106760 | 5106941 | 182 | No | antisense_860 | 5106761 | 417 | RNA |
| as4699 | 5124039 | 5124366 | 328 | No | antisense_870 | 5124354 | 621 | RNA |
| as5028 | 5464169 | 5464418 | 250 | No | antisense_942 | 5464419 | 806 | RNA |
| scr6106 | 6706584 | 6706667 | 84 | No | antisense_1148 | 6706584 | 932 | COD |
| scr6908 | 7672451 | 7672670 | 220 | No | orphan_230 | 7672451 | 916 | RNA |
| scr6925 | 7688336 | 7688466 | 131 | No | orphan_233 | 7688336 | 877 | RNA |

Table S2: Hits of *S. coelicolor*'s predicted ncRNA genes in the `RFAM` database. The **accession code** column corresponds to the identification used within `RFAM`, while the column **ID** refers to the label given to the transcript by `TSS-Captur`. **Start** and **End** refer to the local coordinates within each transcript that were aligned. The **predicted length** refers to the length predicted by `TSS-Captur`, while the aligned length refers to how much of the transcript was actually aligned. Hits marked with an asterisk (*) represent transcripts that come from the data of Vockenhuber *et al.*.

| **ID** | **Target name** | **Acc. code** | **Start** | **End** | ***E*-value** | **Pred. length** | **Aligned length** |
|---|---|---|---|---|---|---|---|
| orphan_87 | 6C | RF01066 | 25 | 100 | $1.70 \times 10^{-}05$ | 167 | 76 |
| antisense_111 | Cobalamin Riboswitch | RF00174 | 229 | 331 | $4.40 \times 10^{-}14$ | 330 | 103 |
| orphan_20 | *Cobalamin Riboswitch | RF00174 | 17 | 235 | $4.90 \times 10^{-}27$ | 153 | 219 |
| orphan_233 | *Scr6925 | RF02826 | 1 | 130 | $2.10 \times 10^{-}25$ | 560 | 130 |
| antisense_192 | *Scr1601 | RF02830 | 6 | 213 | $3.00 \times 10^{-}40$ | 214 | 208 |
| antisense_840 | tRNA | RF00005 | 401 | 468 | $7.10 \times 10^{-}12$ | 467 | 68 |
| orphan_127 | ydaO-yuaA | RF00379 | 182 | 338 | $1.50 \times 10^{-}26$ | 337 | 157 |
| orphan_128 | Scr4115 | RF02829 | 80 | 188 | $2.40 \times 10^{-}16$ | 644 | 109 |
| orphan_157 | Scr5239 | RF02605 | 66 | 224 | $3.90 \times 10^{-}32$ | 543 | 159 |
| orphan_167 | che1 | RF02935 | 370 | 471 | $4.70 \times 10^{-}15$ | 486 | 102 |
| orphan_78 | Bacterial ssrRNA | RF00177 | 602 | 743 | $6.80 \times 10^{-}32$ | 739 | 142 |

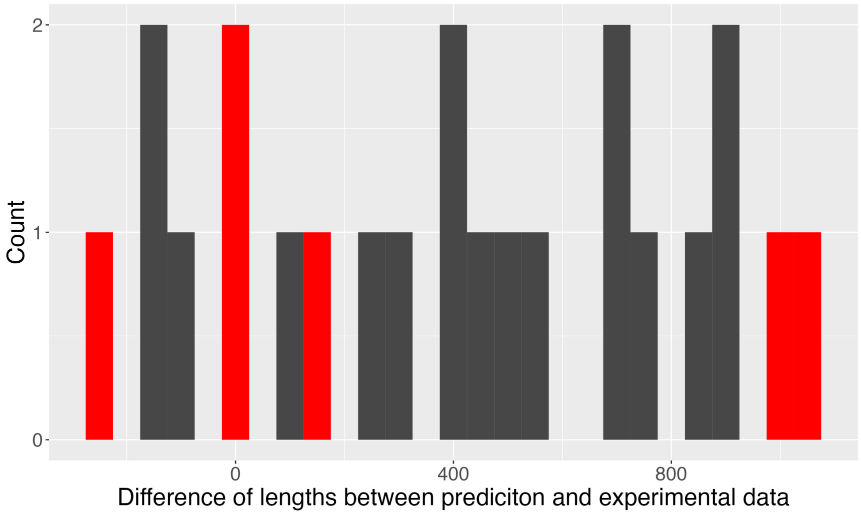


Figure S4: Histogram (bin width = 50) showing the length differences between experimental results presented by Vockenhuber *et al.* [33] and predicted transcripts with `TSS-Captur` for *S. coelicolor*. Highlighted bars (red) represent transcripts confirmed using Northern Blot.
